# BioSAXS measurements reveal that two antimicrobial peptides induce similar molecular changes in Gram-negative and Gram-positive bacteria

**DOI:** 10.1101/611608

**Authors:** A.R. von Gundlach, M. Ashby, J. Gani, P. M. Lopez-Perez, A. Cookson, S. Huws, C. Rumancev, V.M. Garamus, R. Mikut, A. Rosenhahn, K. Hilpert

## Abstract

Two highly active short broad-spectrum AMPs (14D and 69D) with unknown mode of action have been investigated in regards to their effect against the Gram-negative bacteria *E. coli* and the Gram-positive bacteria methicillin-resistant *Staphylococcus aureus* (MRSA). Minimal inhibitory concentration (MIC) measurements using a cell density of 10^8^ cfu/ml resulted in values between 16 and 32 μg/ml. Time kill experiments using 10^8^ cfu/ml revealed complete killing, except for 69D in combination with MRSA, where bacterial load was reduced a million times. Small angle X-ray scattering of biological samples (BioSAXS) at 10^8^ cfu/ml was applied to investigate the ultrastructural changes in *E. coli* and MRSA in response to these two broad-spectrum AMPs. In addition, electron microscopy (EM) was performed to visualize the treated and non-treated bacteria. As expected, the scattering curves generated using BioSAXS show the ultrastructure of the Gram-positive and Gram-negative bacteria to be very different (BioSAXS is not susceptible to the outer shape). After treatment with either peptide, the scattering curves of *E. coli* and MRSA cells are much more alike. This data in conjunction with the EM indicates that ribosomes might be effected by the treatment as well as changes in the nucleoid occurs. Whereas in EM it is notoriously difficult to observe changes for spherical Gram-positives, the BioSAXS results are superior and reveal strongly similar effects for both peptides induced in Gram-positive as well as Gram-negative bacteria. Given the high-throughput possibility and robust statistics BioSAXS can support and speed up mode of action research in AMPs and other antimicrobial compounds, making a contribution towards the development of urgently needed drugs against resistant bacteria.

## Introduction

The WHO has classified antimicrobial resistance as one of the biggest threats to global health and food security. The extent of the threat requires action not only from researchers, but from governments and society. For example, antibiotics are misused in ton-scale in agriculture for growth promotion or prevention of disease, but actions are taken to reduce this, for example the European Union has banned the use of antibiotics for growth promotion in 2006. Currently about 700,000 to 1,000,000 people die worldwide each year because of antibiotic-resistant infections. In the O’Neil Report it is estimated that by 2050 the numbers increase to 10,000,000, more people than are currently killed by cancer (https://amr-review.org/Publications.html). In this report it was estimated that worldwide the additional health care cost for antibiotic-resistant infections will reach US$100 trillion. The situation might even intensify since the number of newly developed antibiotics is steadily declining. FDA approval of new antimicrobials has dropped to three new molecular entities (NME) in this decade.

Antimicrobial peptides (AMPs) are potential novel antimicrobial drugs with some much-desired features, including a low chance of developing drug resistance, fast acting, broad-spectrum activity including multi-drug resistant bacteria. So far only a few have been investigated in clinical studies (Czaplewski et al., 2016; Greber and Dawgul, 2017). There are more than 3,000 natural and artificial peptides described (http://aps.unmc.edu/AP/main.php), the vast majority are cationic. Although they have an enormous variety of sequences and structures, they share certain common features. Cationic antimicrobial peptides are structurally diverse, typically between 5-50 amino acids in length with at least one excess positive charge due to lysine and arginine residues and contain hydrophobic amino acids.

The overall peptide drug market for many different diseases and diagnostics is steadily growing, about 60 peptide drugs approved, with 150 in active clinical trials; and it expected to further grow from US$14.1 billion in 2011 to US$25.4 billion in 2018 (Fosgerau and Hoffmann, 2015; Lau and Dunn, 2018). Demands for peptide drugs has led to A) improved scale up technologies, B) new large-scale GMP certified manufacturing facilities and C) innovative drug administration regimes. These recent developments in peptide drugs has coincided with an increasing cost of novel non-peptide antibiotics, meaning AMPs might soon become a viable economic option for urgently needed new antimicrobial drugs. In the last two decades of AMP research it has become clear that these molecules have multiple biological activities, including antimicrobial, antiparasitic, anticancer and immunomodulatory properties (Mahlapuu et al., 2016). In the same time period multiple bacterial targets of AMPs were discovered (Brogden, 2005), for example binding to RNA, DNA or histones (Cho et al., 2009; Hale and Hancock, 2007; Kobayashi et al., 2000; Xie et al., 2011) blocking DNA-dependent enzymes (Hilpert et al., 2010; Marchand et al., 2006), blocking the synthesis of important outer membrane proteins (Carlsson et al., 1991), binding to the chaperon DnaK and the ribosome (Knappe et al., 2016; Krizsan et al., 2015; Mardirossian et al., 2018b) and lipid 2 (de Leeuw et al., 2010; Schmitt et al., 2010). In addition, the effect of such peptides on blood components was studied (Yu et al., 2015). It is possible to use peptide libraries to screen and optimize antimicrobial peptides (Ashby et al., 2017). With the already large number of antimicrobial peptides and there various modes of actions a method to classify such peptides according to their mode of action would be of high value and BioSAXS was recently used to develop such a classification method (Von Gundlach et al., 2016a)

Small angle X-ray scattering of biological samples (BioSAXS), for example proteins, is a powerful method for the characterization of both ordered and disordered structures in biological samples that provides information about the sizes and shapes ranging from a few kDa to GDa (Chen et al., 2018; Kikhney and Svergun, 2015). In the last decades, X-ray technology has matured to allow the study of protein crystals and proteins in solution down to atomic resolution. The short wavelength of the X-rays (< 1Å) is the key for the success as it enables the probing of small structures. Third generation synchrotron facilities and the advent of diffraction limited, fourth generation storage rings in the near future, provide exceptional brilliance that enables rapid data acquisition (Schroer et al., 2018). In conjunction with the latest generation of single photon counting detectors and autosampler based sample delivery systems, hundreds of samples can be measured per hour (Hajizadeh et al., 2018; Pernot et al., 2018). The novelty of this work is the application of SAXS to reveal structural changes on the length scale between 3 nm – 120 nm within bacteria as consequence of treatment with antimicrobial substances. Subtle structural intracellular rearrangements in the bacteria can accurately be probed across large bacterial populations (hundreds of thousands of bacteria) within seconds (Von Gundlach et al., 2016a). We recently demonstrated that antimicrobial compounds with different modes of action lead to different ultrastructural changes, detectable by BioSAXS (Von Gundlach et al., 2016b). Thus, novel compounds with unknown modes of action can be grouped according to their effect on the bacterial morphology and new responses can be identified.

We have applied different prediction methods for short cationic antimicrobial peptides using artificial neural networks and fuzzy logic, some of which showing great accuracy rates (Cherkasov et al., 2009; Fjell et al., 2009; Mikut et al., 2016). Some of these peptides were tested against *Mycobacterium tuberculosis* and other Gram-positive and Gram-negative microbial pathogens (Ramón-García et al., 2013). We selected two with broad spectrum activity (14L and 69L) for the BioSAXS experiment. In order to achieve proteolytic stability in high bacteria numbers, stereoisomers were used. D-peptide forms (14D and 69D) also showed broad-spectrum activity and the time kill experiments using the D-peptides demonstrated a fast-acting mode of action even with a high number of bacteria present.

## Materials and Methods

### Bacterial strains

Bacterial strains used for antimicrobial activity testing in this project were methicillin-resistant *Staphylococcus aureus* (*S. aureus*) HO 5096 0412 (a neonatal infection isolate, isolated in Ipswich, England in 2005), a methicillin sensitive *Staphylococcus aureus* (ATCC 29213), *Escherichia coli* (*E. coli*, UB1005) F^−^ nalA37 metB1), *E. coli* (ATCC 25922), *E. coli* (68610Y), a clinical isolate from St. George’s University Hospitals NHS Foundation Trust, London, UK, resistant to Gentamicin, Ciprofloxacin and Ceftazidime, obtained from Timothy Planche, wild-type *Salmonella enterica* ssp. Typhimurium (*S. typhimurium*), *Enterococcus faecalis* (*E. faecalis* ATCC 29212), a clinical isolate of *Staphylococcus epidermidis* (*S. epidermidis*) obtained from Dr. Robert E.W. Hancock (Department of Microbiology and Immunology, University of British Columbia) and vancomycin resistant *Enterococcus faecalis* (NCTC 12203).

### Peptides

Antimicrobial peptides were synthesized by automated solid-phase peptide synthesis (SPPS) on a MultiPep RSI peptide synthesizer (Intavis, Tuebingen; Germany) using the 9-fluorenyl-methoxycarbonyl-tert-butyl (Fmoc/tBu) strategy. Reactive side chains were protected by *t*Bu (Tyr and Asp), trityl (Trt, for Asn, Cys, Gln and His), 2,2,4,6,7 pentamethyldihydrobenzofuran-5-sulfonyl (Pbf, for Arg) and *tert*-butoxycarbonyl (Boc, for Lys and Trp). For automated SPPS four equivalents of Fmoc amino acids (Bachem, Bubendorf, Switzerland) were coupled on TentaGel® HL RAM resin (25 μmol scale, loading 0.3-0.4 mmol/g, Rapp Polymere, Tuebingen, Germany) after *in situ* activation with four equivalents of N,N,N′,N′-Tetramethyl-O-(1H-benzotriazol-1-yl)uronium hexafluorophosphate (HBTU; Carbosynth, Berkshire, United Kingdom) and eight equivalents of N-Methylmorpholine (NMM, Sigma, Dorset, United Kingdom). After double coupling procedure (2×30 min) the Fmoc group was cleaved using 20% (*v*/*v*) piperidine (Thermofisher Acros Organics, Geel, Belgium) in dimethylformamide (DMF, Jencons-VWR, Leicestershire, United Kingdom). Peptide amides were cleaved from the resin with 95% (*v*/*v*) aqueous trifluoroacetic acid solution (TFA, Fisher Scientific, Loughborough, United Kingdom) containing 5% (*v*/*v*) triisopropylsilane (TIPS, Thermofisher Acros Organics, Geel, Belgium) / water (1:1) scavenger mixture within 3 h. Cleaved peptides were precipitated from ice-cold methyl *tert*-butyl ether (MTBE; Thermofisher Acros Organics, Geel, Belgium). After washing and collection by centrifugation crude peptides were dissolved in 20% (*v*/*v*) acetonitrile (ACN, Jencons-VWR, Leicestershire, United Kingdom) / 80% (*v*/*v*) water containing 1% (*v*/*v*) TFA to a concentration of 15 mg/ml and analysed by analytical reversed-phase (RP) HPLC on a Shim-pack VP-ODS (120 Å, 150×4.6 mm, Shimadzu, Milton Keynes, United Kingdom) using a Shimadzu LC2010AHT system. The binary solvent system contained 0.1% (*v*/*v*) TFA in H_2_O (solvent A) and 0.1% (*v*/*v*) TFA in acetonitrile (solvent B). The identity was verified by a liquid chromatography electrospray ionisation mass spectrometry (LC-ESI-MS) Shimadzu LC2020 system equipped with a Jupiter 4μ Proteo C18 column (90 Å, 250×4.6 mm, Phenomenex, Cheshire, United Kingdom). The binary solvent system contained 0.01% (*v*/*v*) TFA in H_2_O (solvent A) and 0.01% (*v*/*v*) TFA in acetonitrile (solvent B).

Crude peptides were purified to homogeneity of >92% by preparative RP HPLC on a Shimadzu LC2020 system equipped with a Jupiter 10μ Proteo C18 column (90 Å, 250×21.2 mm, Phenomenex) using a linear gradient system containing 0.01% (*v*/*v*) TFA in H_2_O (solvent A) and 0.01% (*v*/*v*) TFA in acetonitrile (solvent B). Pure products were finally characterized by analytical RP-HPLC and LCMS.

### Bacteriological media and culture conditions

Mueller Hinton broth (MHb) (Merck) was used for all bacterial cultures. Media were prepared and sterilized according to the manufacturer’s’ instructions. Cultures were incubated at 37°C for 18-20 h with aeration and cultures on solid media were incubated at 37°C for 18-24 h.

### Minimal Inhibitory Concentration determination

Minimum inhibitory concentrations (MIC) were determined using a broth microdilution assay as previously described (Wiegand et al., 2008). Bacteria from an overnight culture grown at 37°C were diluted in fresh MHb to achieve a concentration of 1 × 10^6^ CFU/ml. A bacterial suspension (100 μl) was added to wells in a 96 well polypropylene microtiter plate that had been preloaded with serial dilutions of antimicrobial peptides in MHb (100 μl) giving a final bacterial concentration of 5 × 10^5^ CFU/ml. Microtiter plates were incubated at 37°C for 18-20 h before the MIC was determined as the lowest concentration of antimicrobial able to inhibit visible growth.

To determine the MIC towards 10^8^ CFU/ml (MIC_10^8_), which was the bacterial concentration used in BioSAXS experiments, bacteria from an overnight culture were diluted 1:100 in fresh MHb and incubated in a shaking incubator at 37°C and 250 RPM until an OD_600_ of 0.25 was reached, which equated to approximately 2 × 10^8^ logarithmically growing CFU/ml. The MIC was then performed as above without a further dilution of the culture. After 18-20 h incubations, 10 μl of a 500 μM resazurin solution (Sigma-Aldrich) was added to each well of the microtiter plate and the cell viability was determined after a further 1 h incubation by the colorimetric reaction that occurs in the presence of viable cells.

### Time-Kill curves

Overnight cultures of MRSA and *E. coli* UB1005 were diluted 1:100 in MHb and placed in a shaking incubator at 37 ⁰C until an OD_600_ of 0.25 was reached. The culture was then diluted in MHb to achieve a concentration of ~1 × 10^8^ CFU/ml, this culture was split into 1.5 ml tubes. Antimicrobial peptides were added at concentrations of 2 × MIC and sterile water was used as a negative control. Samples were then placed in a shaking incubator set to 37 °C. After 0, 10, 20, 40, 60, and 240 mins, 20 μl of the sample was removed and ten-fold serial dilutions in 10 mM Tris buffer were performed to 10^−6^. From each of the dilutions 5 × 5 μl was plated onto Mueller Hinton agar. Agar plates were placed in a 37⁰C incubator and colony forming units (CFU’s) were counted after 24 h incubation.

### Sample preparation for BioSAXS

A 300 μl aliquot of an overnight culture of MRSA or *E. coli* UB1005 culture was diluted 1:100 in MHb and placed in a shaking incubator set to 37 ⁰C until an OD_600_ of 0.25 was reached. The cultures were then aliquoted into several 2 ml clear plastic vials and doses of peptide were added to achieve a final concentration 2 × MIC_10^8_ (MIC determined for 10^8^ CFU/ml). Additional vials containing culture, but no drug were included as negative controls. The vials were placed in a shaking incubator (250 rpm) at 37°C and incubated for 40 min. Each sample was then washed twice by centrifugation (SciQuip LTD, UK) at 10,000 RPM for 5 mins. Each wash involved the supernatant being removed and pellet resuspended in 1 ml 0.1 M PIPES buffer (pH 7). Samples were then centrifuged for a third time and the pellet was resuspended in 1 ml of 2.5 % glutaraldehyde v/v in PIPES buffer. The samples were then shaken at room temperature for one hour and then washed three times in PBS buffer, at the end of the final washing step the pellet was resuspended in 100 μl of PBS. All samples were then refrigerated at 5°C before analysis.

### Small angle X-ray scattering

The small angle scattering experiments were performed at the BioSAXS beamline P12 at PETRA III (EMBL/DESY) in Hamburg, Germany as in previous experiments. A photon flux of 5 × 10^12^ s^−1^ focused to a spot size of 0.2 mm × 0.1 mm (horizontal × vertical) and the resulting diffraction pattern were recorded with a Pilatus 2M detector (Dectris, Switzerland). The sample (20 μl) was delivered into a cooled glass capillary (20 °C) by an automated sample robot. For each sample 20 diffraction patterns were recorded with an exposure time of 0.05 s. Before and after every sample the background was measured. After angular integration to obtain one-dimensional scattering curves, the background subtraction was performed. To avoid introduction of artefacts by radiation damage, curves collected in subsequent illuminations are compared by a standard F-test (Franke et al., 2012). Only curves collected before the occurrence of radiation damage were further processed. This primary data processing steps were performed using the automated data pipeline SASFLOW.

### Data evaluation

Scattering data was analysed using the open source data mining MATLAB® toolbox Gait-CAD and its successor SciXMiner, using the “Peptide Extension” tool (Mikut, 2010; Mikut et al., 2017). At first, the first data points afflicted by beamstop were removed. To compensate for the experimental variation of the cell density, the data was normalized to the initial region (0.04 nm^−1^ to 0.05 nm^−1^). In order to be consistent with our former data we decided to perform a PCA even with fewer data from these experiments. For the principal component analysis (PCA), the log of the scattering data was used and the range had to be restricted (0.055 nm^−1^ to 0.2869 nm^−1^) due to low intensity of several scattering curves. The principal component analysis is an easy visualization that preserves the main differences of the investigated scattering curves. The sample points are projected to a lower dimensional parameter space, built by so-called principal components. These principal components are orthogonal to each other and remove the redundancies caused by correlations of the sample points. They are computed by finding the eigenvalues of the covariance matrix of the 94 data points per scattering curve. The SAXS data were measured in the q-range of 0.02 nm^−1^ and 4.8 nm^−1^. The 94 data points are contained in the q-range between 0.055 nm^−1^ and 0.2869 nm^−1^ which was used in PCA analysis (Figure 3). The first two principal components were found to describe the variations due to antibiotic treatments. For reasons of better visualization, a centred PCA starting from the mean of all scattering curves *I*_*m*_(*q*) was used: *I*(*q*)=*I*_*m*_(*q*)+*A*·*PC1*(*q*)+*B*·*PC2*(*q*). Consequently, each scattering curve can be approximated by two linear coefficients (*A*, *B*). To provide evidence of reproducibility between two measurements, we measured duplicates of a subset of samples. The mean of the two measurements was used for further analysis. The experimental error estimate given was calculated as average standard deviation of all repeats.

### Electron microscopy

MRSA and *E. coli*, untreated and treated with 14D and 69D for 40 minutes, were subjected to an ethanol series of 30%, 50%, 70%, 95% and three changes of 100% for at least an hour. The samples were transferred to a 1:2 mixture of ethanol to LR White – Hard Grade (London Resin Company, UK) resin then a 2:1 mixture of ethanol to resin and finally 100% resin overnight @ 4°C. The next morning, the resin was removed and replaced with fresh resin and later that day the samples were placed in size 4 gelatine moulds (Agar Scientific), filled with fresh resin and polymerised overnight in an oven at 60°C. 2 μm thick sections were cut which contained the bacteria and these were dried in drops of 10% ethanol on glass microscope slides. They were stained with AMB stain (Azur II & Methylene blue, both Sigma Aldrich Ltd, UK), and photographed using a Leica DM6000B microscope. Ultrathin 60–80 nm sections were then cut on a Reichert-Jung Ultracut E Ultramicrotome with a Diatome Ultra 45 diamond knife and collected on Gilder GS2X0.5 3.05 mm diameter nickel slot grids (Gilder Grids, Grantham, UK) float-coated with Butvar B98 polymer (Agar Scientific) films. All sections were double-stained with uranyl acetate (Agar Scientific) and Reynold’s lead citrate (TAAB Laboratories Equipment Ltd, Aldermaston, UK) and observed using a JEOL JEM1010 transmission electron microscope (JEOL Ltd, Tokyo, Japan) at 80 kV. The resulting images were photographed using Carestream 4489 electron microscope film (Agar Scientific, UK) developed in Kodak D-19 developer for 4 min at 20 °C, fixed, washed and dried according to the manufacturer’s instructions. The negatives were scanned with an Epson Perfection V800 film scanner and converted to positive images.

## Results

For this experiment two broad spectrum short antimicrobial peptides were selected, that were previously described (Hilpert et al., 2005; Ramón-García et al., 2013), see Figure 1.

**Figure. 1:**
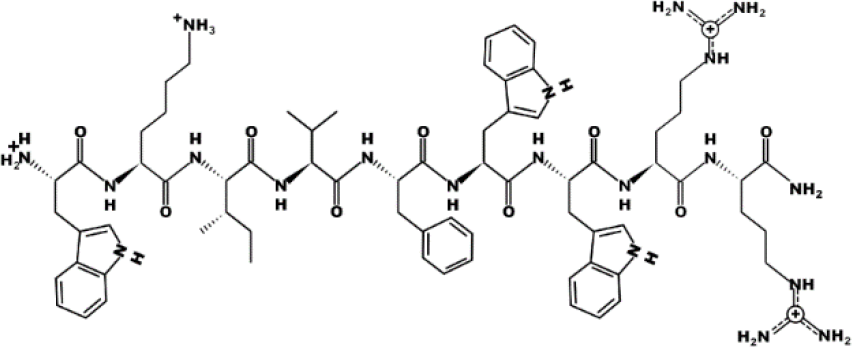
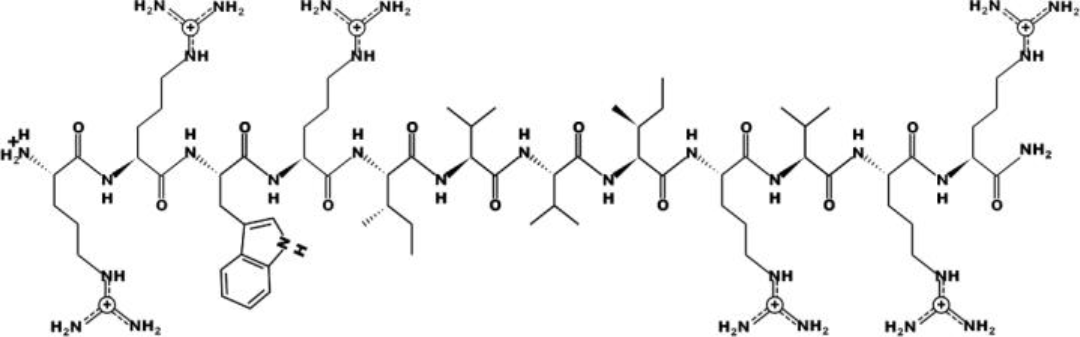
Schematic representation of the peptides 14L (WKIVFWWRR-CONH_2_) (a) and 69L (RRWRIVVIRVRR-CONH_2_) (b)

Peptide 14L (WKIVFWWRR-CONH_2_) is an all-L amino acid peptide comprising of 9 amino acids and peptide 69L (RRWRIVVIRVRR-CONH_2_) is also an all-L amino acid peptide comprising of 12 amino acids. Both peptides are amidated at their C-terminus. MIC values for these peptides against a series of human pathogens are given in Table 1. Since the BioSAXS experiment requires 1,000 times higher bacterial concentrations than are used in classical MIC tests, we decided for this particular experiment to use the all-D peptides in order to avoid problems with fast proteolytic attack by bacterial proteases. These all-D peptides are called 14D and 69D.

**Table 1.** Minimal inhibitory concentration (MIC) in μg/ml for four peptides against several Gram-positive and Gram-negative bacteria. All MICs were performed in Mueller-Hinton bouillon at least three times and data are stated as the modal value. MRSA stands for methicillin-resistant *Staphylococcus aureus* and VRE for vancomycin resistant *Enterococcus faecalis.*

| Peptide/<br>Bacteria | *S.*<br>*aureus*<br>ATCC<br>29213 | MRSA | *E. coli*<br>ATCC<br>25922 | *E. coli*<br>UB1005 | *E. coli*<br>68610Y | VRE<br>NCTC<br>12203 | *E.*<br>*faecalis*<br>ATCC<br>29212 | *S.*<br>*epidermidis* |
| --- | --- | --- | --- | --- | --- | --- | --- | --- |
| T14L | 0.5 | 0.5 | 2 | 4 | 2 | 1 | 1 | <0.25 |
| T14D | 1 | 1 | 2 | 2 | 2 | 1 | 1 | <0.25 |
| T69L | 2 | 2 | 2 | 2 | 2 | 2 | 4 | 0.5 |
| T69D | 2 | 1 | 2 | 2 | 2 | 2 | 2 | 0.5 |

For a MIC determination, an inoculum of about 2-5×10^5^ bacterial cells is used, however for the BioSAXS a bacterial density of about 1×10^8^ cells is required and in consequence more peptide molecules are needed in order to kill or inhibit bacterial growth. The MIC for both peptides with an inoculum size of 10^8^ was determined with 26 μg/ml for 14D and 32 μg/ml for 69D against *E. coli*, and 16 μg/ml for 14D and 32 μg/ml 69D against MRSA. For the time kill assay twice the MIC_10^8_ concentration was used. Both the peptides were able to kill 1×10^8^ bacterial cells completely, except for 69D against MRSA, were the bacterial load was reduced a million times, see Figure 2.

**Figure 2:**
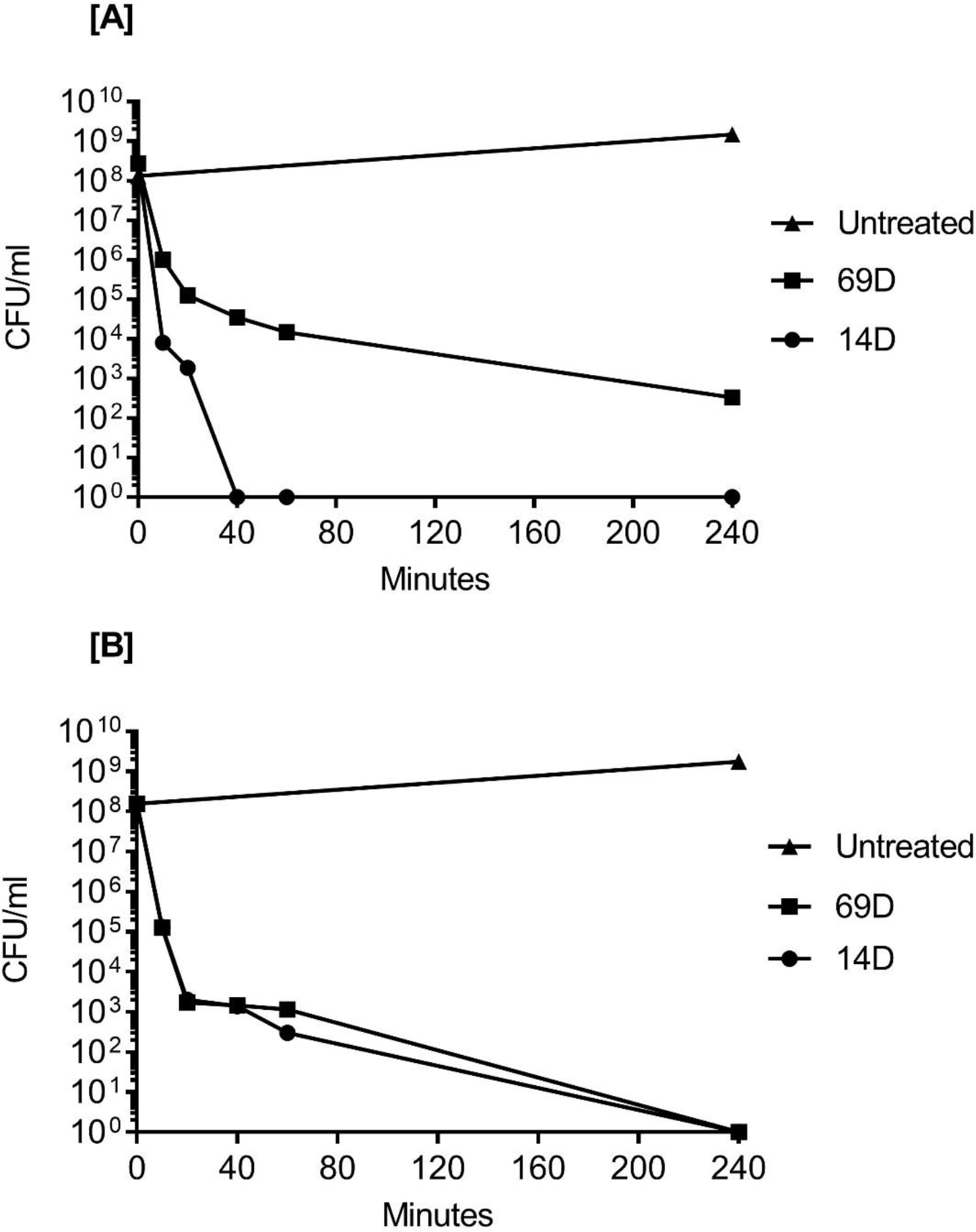
Time kill curves of [A] MRSA and [B] *E. coli* following incubation with the antimicrobial peptides 14D and 69D in Mueller Hinton broth. Peptides were used at twice the MIC required to inhibit the growth of 1 × 10^8^ CFU/ml for each organism.

Based on the results of the time kill assay the time points 10 and 40 mins were chosen for the BioSAXS measurements. Using two times the MIC_10^8_ (required to inhibit the growth of 1 × 10^8^ CFU/ml) and 10^8^ cells, peptides 14D and 69D were incubated with the bacteria and after 10 and 40 minutes samples were taken to be processed for BioSAXS measurement and at 40 min for electron microscopy (see Material and Methods). The results of the scattering are shown in Figure 3.

**Figure 3.**
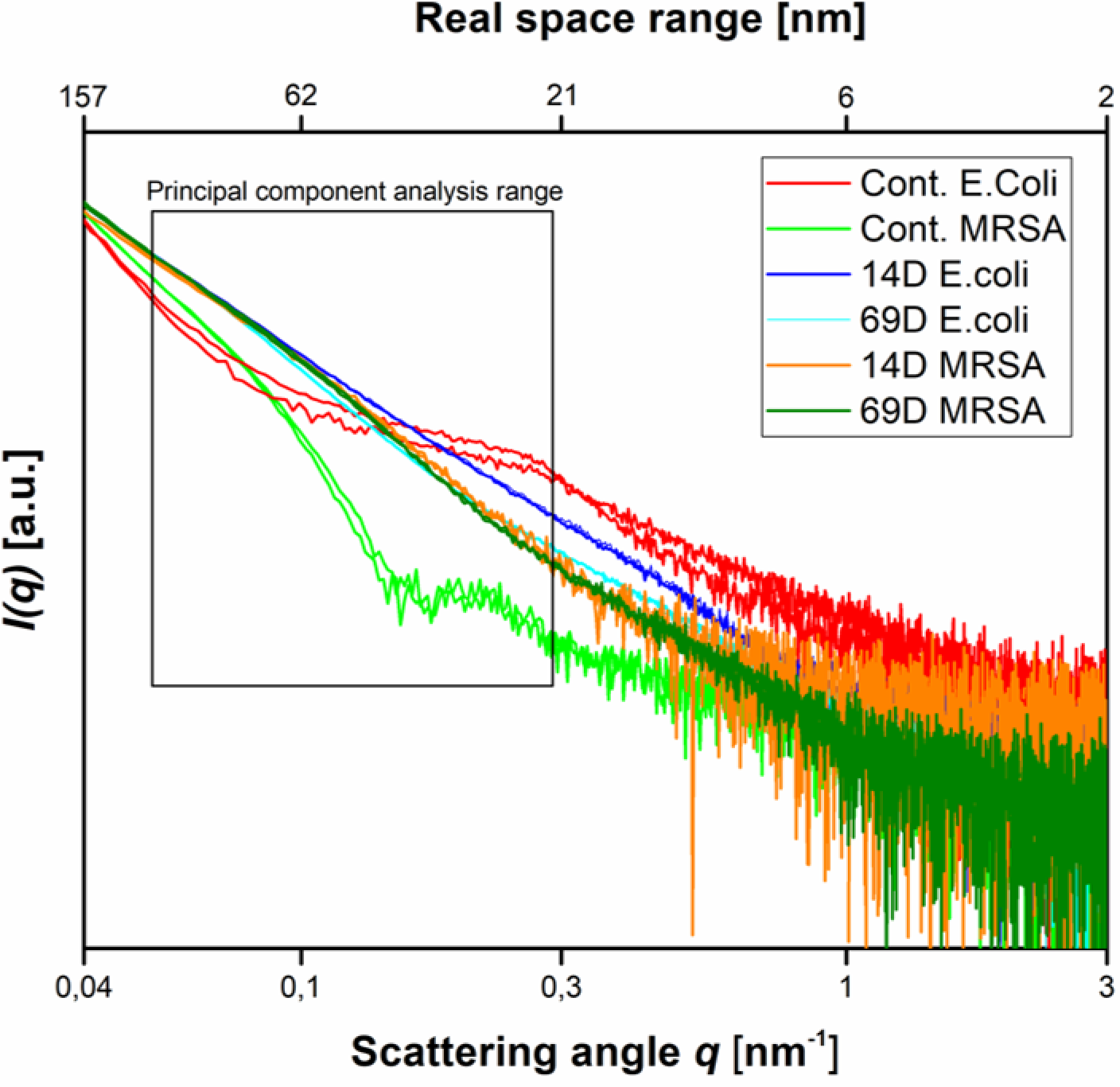
Scattering data as measured at the P12 BioSAXS beamline at PETRA III (Hamburg, Germany) at a photon energy of 8 keV. Scattering data from *E. coli*, MRSA untreated and treated with peptides 14D and 69D at 40 min, measured in duplicate (shown as separate curves). The box indicates the range that was used to calculate the principal component analysis.

With respect to the size range covered by the small angle X-ray scattering experiments, untreated cells of *E. coli* and MRSA differ mainly between a size range of 20 nm to 60 nm, with higher contribution from the Gram-negative cell. The measurement is only susceptible to the internal structure and not the outer shape of the bacteria. After treatment with either peptide, the scattering curves of *E. coli* and MRSA cells are much more alike – smoother with a constant slope. In order to better visualize the differences in the scattering curves, a principle component analysis was performed using the curve section shown in Figure 3. The result of this analysis and the electron microscopy images of *E. coli* and MRSA at 40 min are presented in Figure 4.

**Figure 4:**
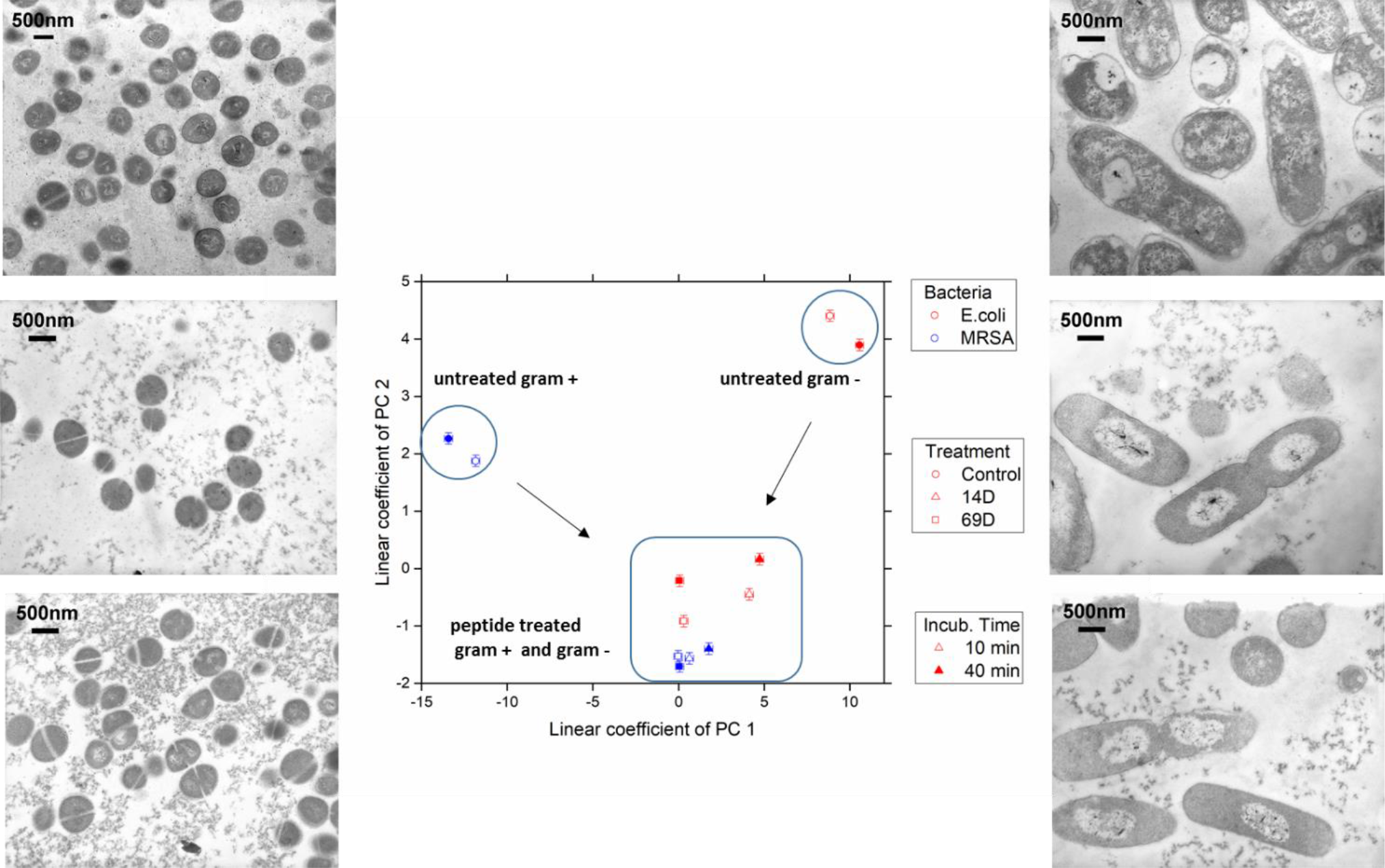
The linear coefficients of the first two principle components discriminate morphological changes and modes of action. The colour decodes the bacterial species, the symbol the applied treatment and the symbol thickness the incubation time. The error estimate is calculated from duplicate measurements and found to be 0.24 for the coefficient of PC1 and 0.10 for the coefficient of PC2. The transmission electron micrographs at a 15,000 times magnification show MRSA left and *E. coli* right hand side. Top row shows untreated cells, middle row treatment with 14D and lower row treatment with 69D, peptide treatment time: 40min.

## Discussion

Antimicrobial peptides are potential novel antimicrobial drugs with some much-desired features, for example a low chance for the development of resistance, fast acting, broad spectrum of activity and activity against multidrug resistant bacteria. There is a large variety of structures and sequences of AMPs and in recent years it has become clear that there are also a variety of targets (Le et al., 2017). Today, there is a huge body of literature regarding AMPs, however there is only few articles published on target validation, pharmacological and safety studies (Czaplewski et al., 2016; Greber and Dawgul, 2017; Mardirossian et al., 2018a; Schmitt et al., 2010). This contributes to the fact that only a few AMPs are enrolled in clinical studies. We have already shown that BioSAXS can support research on antimicrobials to select compounds with possible new modes of action and therefore select compounds with alternative mode compared with mechanisms of action of failing conventional antibiotics (Von Gundlach et al., 2016a). In this study we compared the effects on Gram-positive and Gram-negative bacteria to further understand the broad-spectrum activity of two antimicrobial peptides.

The effect of an antimicrobial compound on bacteria can be quite complex. The compound will act on their target(s) and induces changes at this site which can lead to secondary effects at the same, or at different sites. In case the target(s) are inside the bacteria the compounds will cross the outer envelopment and the membrane and could therefore cause additional changes. At the same time, the bacteria reacts to the compound and induce several stress responses and coping mechanism in order to survive. The observed effect is consequencently a mixture of ultrastructural changes on the bacterial level caused by direct and indirect effects of the antimicrobial compound as well as direct and indirect effects of the stress response of the bacteria. For each compound these effects will be concentration and time dependent (Von Gundlach et al., 2016a).

The BioSAXS measurements requires a high bacterial density and therefore higher amounts of proteases are present as compared to a conventional MIC test. The proteases could cleave the L-peptides into many different fragments which may render inactive or also interact with the bacteria and prompt a detectable alteration in ultrastructure. To restrict this, the L-peptide sequences were synthesized as complete D-versions that will be extremely stable in the presence of the proteases for the time frame of the experiment. For the BioSAXS experiment only the complete D-versions were used, therefore MIC values and time kill assays were performed using complete D-peptides.

The electron microscopy images show that the treatment of *E. coli* with either peptide results in a separation of the cytoplasm and the nucleoid, which appears to be in the centre of the cell. In addition, the cytoplasm becomes much more homogenous as compared to the control. Interestingly, the peptides in this study result in a very different response compared to a peptide (RLKRWWKFL) described in our previous studies, indicating different modes of action (Von Gundlach et al., 2016a). In addition, we could not detect any similarities, for example damages to the cell wall or membrane, dramatic changes in the inside of the cells, typically seen with polymyxin B, a cyclic lipopeptide with detergent-like mode of action (Von Gundlach et al., 2016a). The type of nucleoid separation observed after a treatment with peptide 14D and 69D are similar to the ribosome acting drugs such as chloramphenicol or tetracycline which may indicate a similar target or cell response (Von Gundlach et al., 2016a). From studies on living cells, it is known that an inhibition of the peptide synthesis leads to a compaction of the bacterial nucleoid while an inhibition of the RNA synthesis by rifampicin expands the bacterial nucleoid (Chai et al., 2014). The mechanism for the condensation of the nucleoid is described as the absence of “transertion”, the synthesis of membrane proteins in close proximity to the cell wall. When the protein synthesis is inhibited, the DNA / RNA complexes are no longer tethered to the cell wall which leads to a collapse of the nucleoid in the cell centre (Cabrera et al., 2009). Due to the spherical structure and high cell wall density of *S. aureus*, changes in the electron microscopy images between treated and untreated cells are harder to detect, although treatment with the peptides does also seem to lead to a more homogenous cytoplasm.

The scattering curves generated using BioSAXS show the ultrastructure of the Gram-positive and Gram-negative bacteria to be very different as expected. However, following treatment with both antimicrobial peptides 14D and 69D, the ultrastructure of the MRSA and the *E. coli* became more similar to each other. Drastic change occurs in the range of 20 to 45 nm. The average protein is between 2-10nm (large proteins like IgG about 10 nm), large protein complexes like ribosomes are about 20-30nm and compacted protein/DNA complexes are about 30nm. This data in conjunction with the EM also indicates that ribosomes can be effected by the treatment as well as changes in the nucleoid. The ultracellular effect of 14D and 69D on *E. coli* is similar direction although not the same. The SAXS data reveal a structural difference in the first principle component. While both feature a condensed nucleoid, 69D also seems to affect the cellular wall. In MRSA, both 14D and 69D initiate very similar changes even after 10 min, which remain unchanged after 40 min. In *E. coli* the strong morphological effect of the peptide already comes into play after 10 min. After 40 min the alteration does not increase, rather reaching an equilibrium state.

In conclusion, so far we had only shown that BioSAXS can be used as a method to study effects of antimicrobials on Gram-negative bacteria, here for the first time we show that Gram-positive bacteria can also be used to detect changes after peptide treatment. Whereas in EM it is notoriously difficult to observe changes for spherical Gram-positives, the BioSAXS results are superior and reveal strongly similar effects for both peptides induced in Gram-positive as well as Gram-negative bacteria. Given the high-throughput possibility and robust statistics we believe that BioSAXS can support and speed up mode of action research in AMPs and other antimicrobial compounds, making a contribution towards the development of urgently needed drugs against MDR bacteria.

## Author Contributions

ARvG, CR and VMG performed the BioSAXS measurement. ARvG, CR and RM analysed the data. RM provided a program for data handling. MA, JG, PLP performed MIC studies, time kill experiments and peptide synthesis, purification and characterization. AC performed the transmission electron microscopy, SH supported this work. KH, AR and SH contributed conception and design of the study. ARvG and KH wrote the manuscript.

## Acknowledgments

The excellent support of Dr Clement Blanchet (EMBL) and the staff of PETRA III during SAXS measurements is gratefully acknowledged. KH thanks life for the opportunity to continue on and despite the odds to be able to continue researching.

